# A physiologically-based model of bile acid metabolism in humans

**DOI:** 10.1101/2025.07.19.665677

**Authors:** Bastian Kister, Lars M. Blank, Maike Pollmanns, Theresa H. Wirtz, Lars Kuepfer

## Abstract

Altered serum BA profiles have been reported for several chronic liver diseases. However, relevant modifications in bile acid disposition are difficult to investigate mechanistically due to the interference of the various physiological processes involved at various levels of bile acid metabolism. In this regard, validated computational models represent a promising tool to understand the functional effect of disease-related alterations on bile acid composition along the gut-liver-axis through mechanistic analyses of the underlying physiological processes. We applied a physiologically-based models of bile acid metabolism in humans comprising the five predominant bile acids. The model describes various physiological aspects of bile acid metabolism including synthesis, enterohepatic circulation, hepatic and microbial conversions, postprandial gall bladder emptying as well as excretion via feces and urine. Model development was based on 39 previously published studies in patients without any liver diseases. In the future, the established model may serve as reference for analysis of the role of bile acids in health and disease.

## Introduction

Bile acids (BAs) play vital roles in numerous physiological processes, such as nutrient digestion and hormone metabolism ^1,2^. The BA pool is a complex mix of various BA species. Primary BAs are synthesized from cholesterol in the liver and involves enzymes located in different cellular compartments. Microbial transformation of primary bile acids produces various secondary BAs and primarily occurs in the distal portion of the small intestine and large intestine through processes like deconjugation, dehydrogenation, dehydroxylation, and epimerization ^3,5^. Both primary and secondary BAs can be conjugated with either glycine or taurine in hepatocytes.

In the body, BAs undergo continuous enterohepatic circulation (EHC) between the liver and the intestine. The liver secretes BAs into the bile canaliculi, where they accumulate in the gallbladder. Upon food intake, gallbladder contractions release significant amounts of stored BAs into the small intestine to aid in lipid absorption. Enterocytes actively reabsorb most BAs, primarily in the ileum, and release them into the portal blood. The remaining BAs are either absorbed through passive diffusion or excreted in the feces. BAs from the portal blood are efficiently reabsorbed into the liver, potentially reaching vascular circulation and other tissues through sinusoidal transport.

Owing to the systemic nature of BA metabolism, diseases affecting both the liver (e.g. liver cirrhosis, liver cancer, or inflammatory bowel disease) and the intestine (e.g., ulcerative colitis or Crohn’s Disease) have been linked to modifications in BA composition and distribution ^2,7-14^. Investigating such alterations, however, can be challenging due to the intricate physiological processes involved and the need for invasive sampling techniques.

In this work, we developed a physiologically-based model of human BA metabolism at the whole-body level which may be used as a platform for mechanistic investigation of BA metabolism. Our model describes the physiology of human BA metabolism in detail and can be used to simulate tissue concentration profiles of the most abundant BAs in human. To inform and validate the model, a comprehensive data set was compiled from literature. The model may be used as a tool for hypothesis testing and as a bridge between discoveries within mouse studies and clinical applications in human patients.

## Results

The computational framework for modeling human BA metabolism is built upon a physiologically intricate representation of the organism, encompassing various processes such as synthesis, hepatic and microbial transformations, systemic and enterohepatic circulation, and the elimination of BAs from the system. Figure 1 provides a visual overview of the model’s structure.

**Figure 1.**
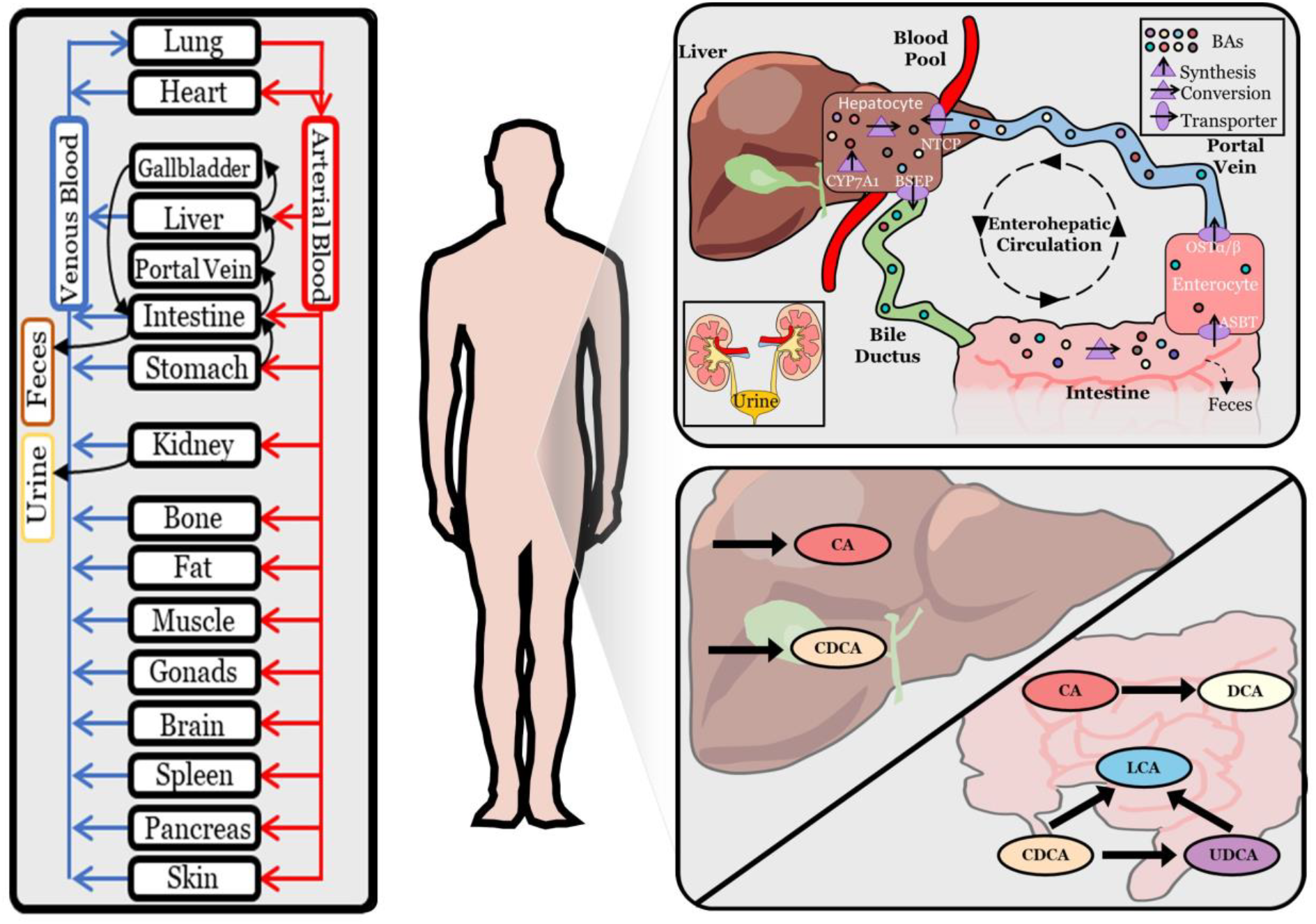
Physiologically-based bile acid model. Schematic overview of a physiologically-based model of BA biosynthesis via CYP7A1, hepatic and microbial transformation, active transport processes via BSEP, ASBT, OSTα/β and NTCP, as well as fecal and renal excretion. Reactions of BAs are located either in the intracellular space of the liver or in the intestinal lumen.

The development of this model employed a methodology rooted in physiologically-based pharmacokinetic (PBPK) modeling, as detailed in Kuepfer et al. ^15^. Within the PBPK framework, the metabolites of BAs are depicted as molecules circulating in the system. This modeling approach offers a comprehensive portrayal of human physiology by integrating specific information on organ volumes, tissue composition, organ surface areas, and blood perfusion rates, all grounded in prior knowledge ^16,17^.

A notable aspect of the mechanistic structure of the PBPK model is its capacity to extend its applicability to novel situations and conditions. This adaptability enhances its utility in representing diverse scenarios within the intricate landscape of human BA metabolism.

To determine the kinetic parameters of the model, an extensive dataset is essential. A comprehensive literature review was hence undertaken, collating BA measurements from 39 studies as detailed in Table 1. This dataset encompassed diverse baseline measurements in venous and portal blood, liver, feces, and bile. Additionally, it included postprandial responses in blood, as well as data on synthesis, excretion, and flow rates.

**Table 1.** Overview of the literature data set. Shown are the number of studies that could be found reporting bile acid measurements in specific organs or for processes. See also supplementary tables 1-8.

| Organ | Number of studies |
| --- | --- |
| Systemic circulation | 18 |
| Portal blood | 4 |
| Liver | 4 |
| Synthesis rate | 5 |
| Renal excretion | 4 |
| Feces | 2 |
| *Bile* | *1* |
| *Intestinal bile flow* | *2* |
| *Single meal studies* | *5* |
| *Multiple meal studies* | *2* |

To address variability among measurements from different studies, effect sizes were synthesized using a meta-analysis approach utilizing a Bayesian mixed-effect model with weakly informative priors ^18-20^.

It is crucial to acknowledge that while this dataset forms a robust basis for model development, there exist qualitative and quantitative variations among the diverse data points defining this ‘healthy’ reference state. The statistical foundation for BA levels, particularly in systemic circulation, is robust and encompasses all pertinent BA species. However, information from other organs relies on fewer studies and individuals, providing only partial coverage of the BA pool.

The model considers four fundamental transportation mechanisms:

1. BSEP facilitates the export of BAs from the liver to either bile or the duodenum;
2. ASBT is responsible for the uptake of BAs from the intestinal lumen;
3. Enterocytes release BAs into the portal bloodstream using OSTα/β;
4. NTCP facilitates the absorption of BAs from the portal bloodstream into hepatocytes.

No data on the expression of ASBT and OSTα/β along the gut segments was available. Thus, it was assumed that ASBT is uniformly expressed in the small intestine, with no expression in the large intestine, while OSTα/β is consistently expressed in both small and large intestine at a constant level, but the latter with higher expression levels.

Moreover, postprandial responses in BA levels can be observed in humans, and several studies measuring BA concentration after a meal are accessible. Therefore, postprandial dynamics and food intake were integrated into the models, considering varying numbers of meals per day and different intervals between meals. For model establishment, three meals per day with 4-hour breaks between meals (0h-4h-8h) were considered. This was facilitated by the modeling software used, which incorporates a function triggering gallbladder emptying into the duodenal lumen upon food intake. Parameters governing the kinetics of this process had to be adjusted based on experimental data but were confined within physiologically reasonable ranges.

In addition to incorporating physiological aspects of the organism, essential components of PBPK models also encompass physicochemical properties such as molecular weight, solubility, lipophilicity, and binding to plasma proteins ^15^. These features play a critical role in determining distribution between organs and plasma, as well as in passive transport, achieved through suitable distribution models. In this context, the physicochemical properties of glyco-conjugated forms (Table 2) were utilized to define the compound attributes within the PBPK model for small molecules.

**Table 2.** Physicochemical properties of bile acids used.} Overview of physicochemical properties and their source that were used to inform compound specific parameters in the PBPK model. For the total BA (total CA: tCA, total CDCA: tCDCA, total UDCA: tUDCA, total DCA: tDCA, total LCA: tLCA) the corresponding physicochemical values of the glyco-conjugated form (G-BA) were taken.

| *BA species* | *Property* | *Value* | *Source* |
| --- | --- | --- | --- |
| tCA | MW [g/mol] | 465.6 | PubChem CID 6675 |
| tCA | Solubility [g/l] | 0.025 | ALOGPS (HMDB<sup>21</sup>) |
| tCA | logP | 0.07 | Heuman et al.<sup>22</sup> |
| tCA | pKa (acidic) | 3.77 | ChemAxon (HMDB<sup>21</sup>) |
| tCA | pKa (basic) | -0.17 | ChemAxon (HMDB<sup>21</sup>) |
| tCA | FU | 0.3247 | Predicted<sup>23</sup> |
| tCDCA | MW [g/mol] | 449.6 | PubChem CID 387316 |
| tCDCA | Solubility [g/l] | 0.0079 | ALOGPS (HMDB <sup>21</sup>) |
| tCDCA | logP | 0.51 | Heuman et al. <sup>22</sup> |
| tCDCA | pKa (acidic) | 3.77 | ChemAxon (HMDB <sup>21</sup>) |
| tCDCA | pKa (basic) | -0.29 | ChemAxon (HMDB <sup>21</sup>) |
| tCDCA | FU | 0.0966 | Predicted <sup>23</sup> |
| tUDCA | MW [g/mol] | 449.6 | PubChem CID |
| tUDCA | Solubility [g/l] | 0.00135 | ALOGPS (HMDB <sup>21</sup>) |
| tUDCA | logP | -0.89 | Heuman et al. <sup>22</sup> |
| tUDCA | pKa (acidic) | 3.77 | ChemAxon (HMDB <sup>21</sup>) |
| tUDCA | pKa (basic) | -0.49 | ChemAxon (HMDB <sup>21</sup>) |
| tUDCA | FU | 0.0966 | Predicted <sup>23</sup> |
| tDCA | MW [g/mol] | 449.6 | PubChem CID 2733768 |
| tDCA | Solubility [g/l] | 0.072 | ALOGPS (HMDB <sup>21</sup>) |
| tDCA | logP | 0.65 | Heuman et al. <sup>22</sup> |
| tDCA | pKa (acidic) | 3.29 | ChemAxon (HMDB <sup>21</sup>) |
| tDCA | pKa (basic) | -0.61 | ChemAxon (HMDB <sup>21</sup>) |
| tDCA | FU | 0.0965 | Predicted (HMDB <sup>21</sup>) |
| tLCA | MW [g/mol] | 433.6 | PubChem CID 439763 |
| tLCA | Solubility [g/l] | 0.0015 | ALOGPS (HMDB <sup>21</sup>) |
| tLCA | logP | 1.05 | Heuman et al. <sup>22</sup> |
| tLCA | pKa (acidic) | 3.77 | ChemAxon (HMDB <sup>21</sup>) |
| tLCA | pKa (basic) | -1.1 | ChemAxon (HMDB <sup>21</sup>) |
| tLCA | FU | 0.0682 | Predicted (HMDB <sup>21</sup>) |

Bile acid synthesis was modeled as a constant formation rate within the intracellular compartment of the liver. The determination of this synthesis rate and the rate of renal excretion was informed by available literature data (Supplementary Table 4). While explicit data on the rate of fecal excretion was not provided, the model maintained equilibrium over an extended period. Consequently, the cumulative rates of excretion were balanced against the sum of all synthesis rates. These excretion processes were modeled using either passive transport or active clearance, depending on the specific process.

To achieve system stability, the model required simulation spanning several hundred hours and numerous meal events. Furthermore, the model incorporated additional transformations within the liver and by microorganisms, as illustrated in Figure 1. These included reactions involving the dehydroxylation of CA to DCA, and CDCA and UDCA to LCA, as well as the epimerization of CDCA to UDCA.

The overall microbial activity was linked to the estimated bacterial density according to Gorkiewicz et al. ^24^.

### Model calibration and validation to literature data

The model underwent calibration using BA measurements gathered from the literature. This calibration enabled the model to reasonably reproduce BA levels, as depicted in Figure 2. A comprehensive list of final model parameters is provided in Supplementary Tables 9-12. Except for one fasting state/baseline measurement (Figure 2A), all predictions fell within a five-fold range, accounting for the inherent variation in the data. Conversely, the model consistently underpredicted the postprandial flow of total BAs within the small intestinal lumen, although the dynamics followed a similar trend. This was deemed acceptable as the corresponding data was obtained from a single individual in the 1970s ^25^, lacking replicates and raising concerns about reliability.

**Figure 2.**
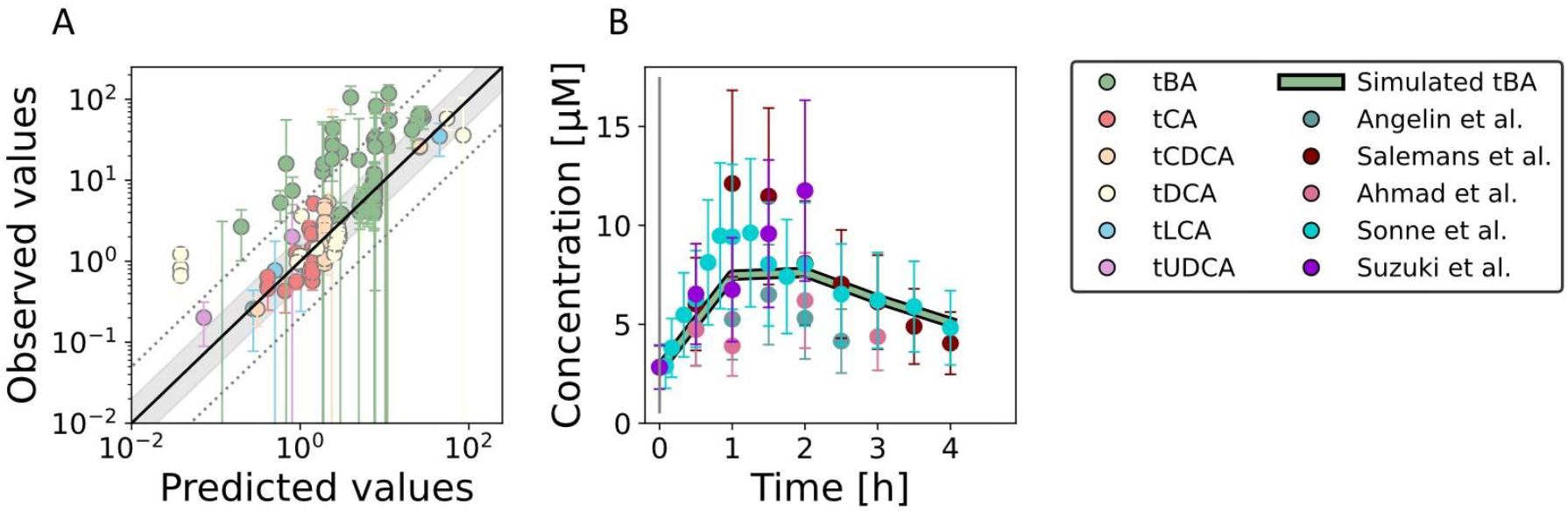
BA model fit to literature data. Model simulations of bile acid concentration in human against corresponding data points used for fitting. Shown are concentration in a fasting state against data points synthesized by meta-analysis of various studies published in literature (A) and postprandial total BA responses of tBA (B) in venous blood plasma. Error bars show the SD. In panel A, the black line represents unity, the gray area the two-fold range and the dashed lines the five-fold range.

In contrast, the dynamics of gallbladder emptying and postprandial BA levels in blood plasma (Figure 2B) were well-captured. The magnitude and dynamics of postprandial responses were accurately predicted for total BAs, total cholic acid, and total deoxycholic acid. However, the response for total chenodeoxycholic acid was generally underpredicted.

Despite the inherent complexities of the modeled system and the scarcity of data, especially outside the bloodstream, the model demonstrated robust agreement between experimental data and simulation outcomes.

To examine the model’s behavior, a sensitivity analysis was performed to discern the influence of the fitted parameters on various BA concentrations derived from the literature (Supplementary Tables 12-16}). Notably, all evaluated BA levels, particularly those of secondary BAs, demonstrated sensitivity to alterations in intestinal pH.

Modifications in BA synthesis, BSEP activity, and plasma protein concentration exerted substantial effects on BA concentrations in venous and portal blood plasma, as well as in the liver. Changes in the functional liver volume predominantly impacted BAs in the liver and venous blood plasma, with the latter also exhibiting sensitivity to NTCP activity. The composition of fecal BAs was most profoundly influenced by parameters associated with microbial activity, ASBT transporter activity, and intestinal transit time.

The accuracy and reliability of any computational model depend on its capacity to replicate real-world observations and experimental data. In this regard, the model underwent testing against literature data that was withheld during model calibration, including postprandial and fasting tBA flow in the duodenum ^26,27^ and postprandial responses in blood plasma from multiple meal studies ^28,29^. The comparison between simulated outcomes and actual measurements, not used in the calibration process, is illustrated in Figure 3.

**Figure 3.**
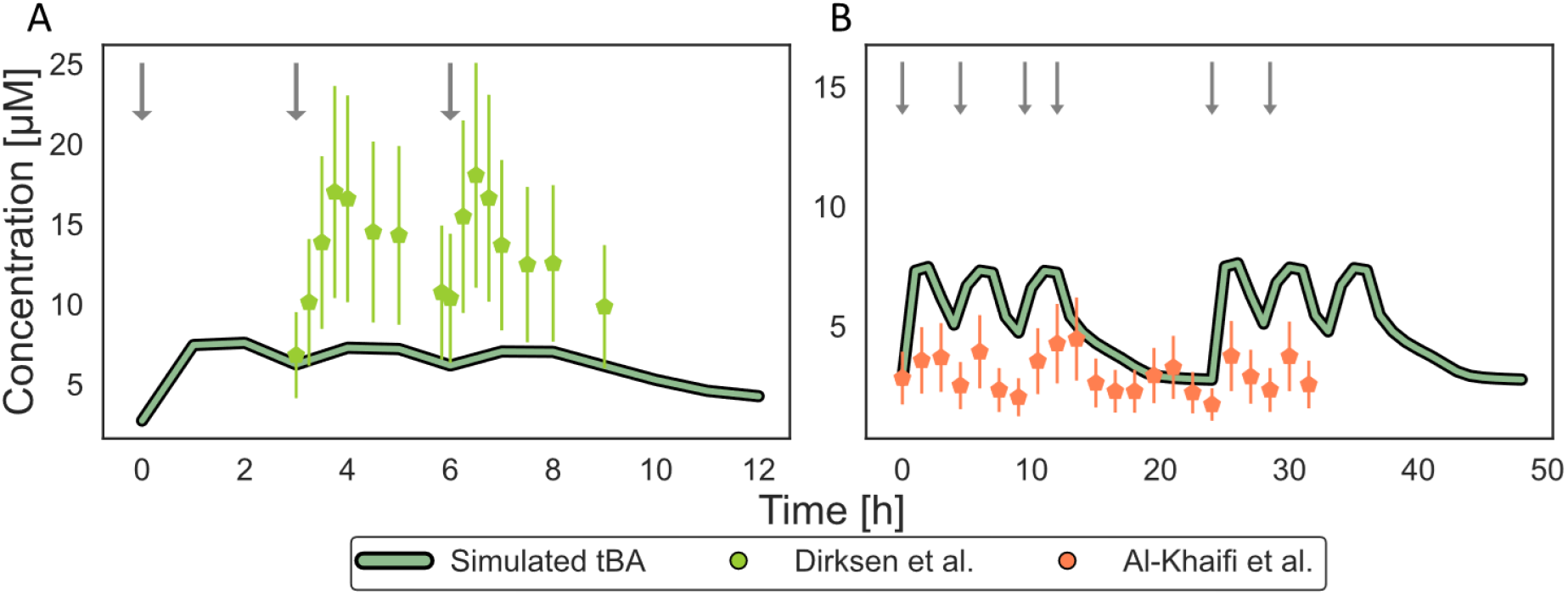
BA model validation to literature data. Model predictions of postprandial tBA concentration in venous blood plasma (A and B) in human against corresponding data points from literature. The model was simulated with 3 meals per day with 3 h between meals (0-3-6) to recapitulate the data from Dirksen et al. ^29^ (A) or with 4 meals per day at 0-4.5-9.5-12 h for data from Al-Khaifi et al. ^28^ (B). Error bars show the SD and arrows indicate a meal event.

Similar to observations during model calibration, predictions of duodenal tBA flow were underestimated. When recapitulating postprandial responses after a second and third meal (Figure 3A), the BA levels were slightly lower than observed, while responses after six meals in 1.5 days were higher (Figure 3B). This outcome was anticipated, as both of these studies reported BA levels that disagreed with the remaining studies utilized in this work (Supplementary Figure 1). While the absolute values of these studies cannot be reconciled when considering the data used for calibration, the general trend was accurately captured.

To enhance the comprehensiveness of model validation, a population simulation involving a virtual cohort of 1,000 healthy individuals was conducted using PK-Sim. The studies by Baier et al. ^30,31^ had previously underscored the necessity for population simulations to adequately address the observed variability in the data.

This cohort featured diverse anthropometric attributes, covering ages ranging from 20 to 60 years and maintaining an equal distribution between sexes (50% females). Body mass index (BMI) within the cohort spanned from 19 to 25 kg/m2. In this simulation context, reference concentrations for all relevant transporters, enzymes, and gallbladder emptying kinetics were varied within a 10% range.

Emphasis was placed on postprandial responses of BAs in blood, considering the wealth of available data demonstrating substantial variability. The results are depicted in Figure 4. Consistent with prior findings, population-level total BA flows in the small intestine were underestimated. In contrast, variability in total BA levels in blood plasma after a meal was accurately captured for both single (Figure 4A) and multiple meals (Figure 4B, C).

**Figure 4.**
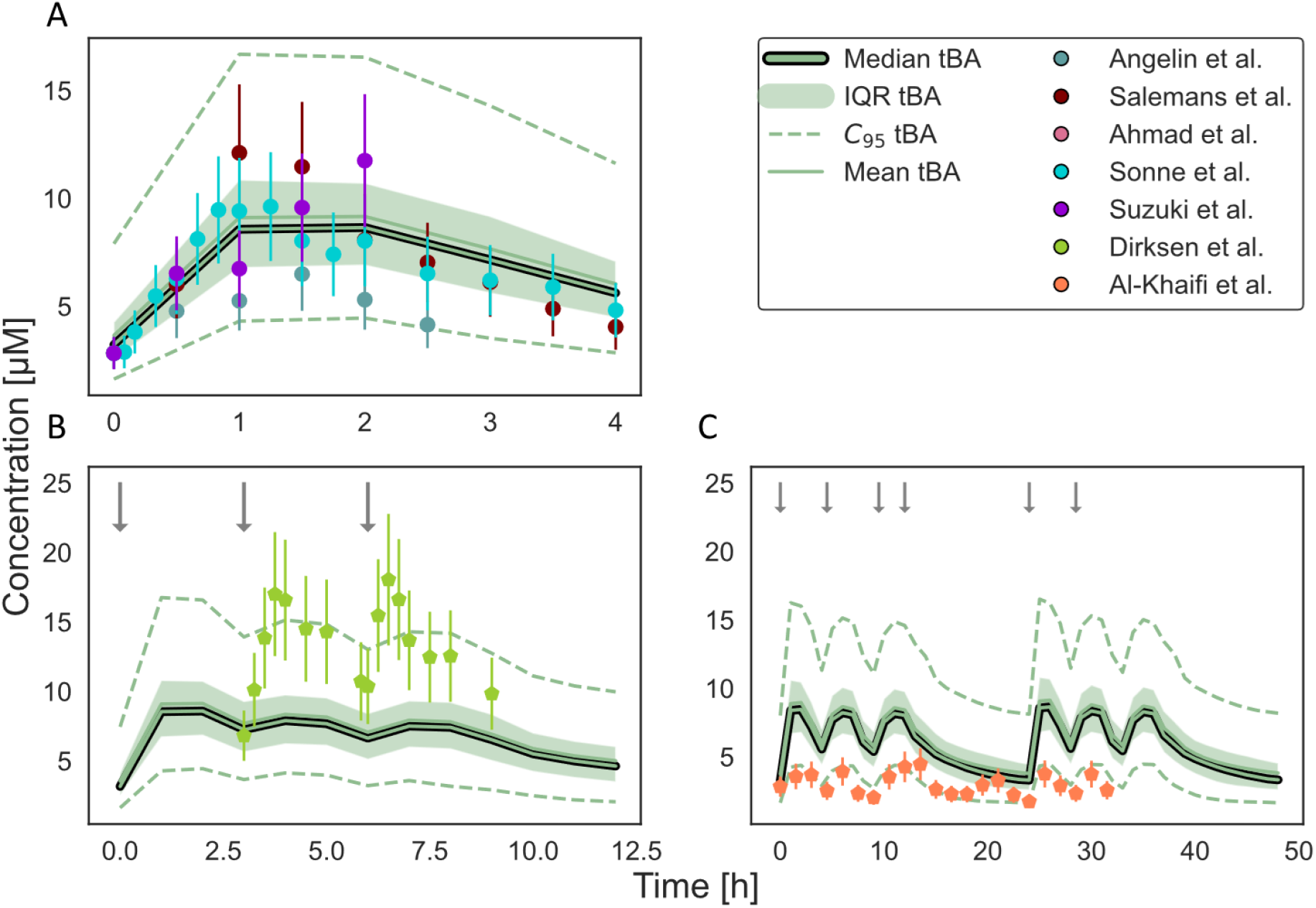
Population simulation of the BA model for the prediction of postprandial responses. Model simulation of postprandial responses of tBA in venous blood plasma in a virtual population of 1,000 individuals after a single meal (A), three meals (B) and six meals (C). Shown are the median (thick line), mean (thin line), IQR (area) as well as the 2.5 and 97.5 percentile (dashed lines) of the population simulations. Error bars show the SD and arrows indicate a meal event.

Despite grappling with the intricacies inherent in the modeled system, the limited availability of data, and the challenge of reconciling conflicting data points, the model demonstrated a commendable level of agreement between experimental data and simulation outcomes.

## Discussion

This study was centered on the construction of physiologically-based models capturing the intricacies of BA metabolism in humans. Due to technical constraints, a specialized model was developed, placing emphasis on quantifying the levels of the five predominant BAs: CA, CDCA, DCA, LCA, and UDCA. The model encompasses critical processes such as systemic circulation, synthesis, hepatic and microbial conversions, gallbladder emptying, and excretion via feces and urine.

The model, carefully established and validated using a dataset from 39 studies characterizing the “healthy” reference, encountered challenges due to data scarcity and reconciling different study findings. Limited information on BA levels beyond postprandial responses in blood plasma within a four-hour time frame post-meal was available.

Concerns about statistical robustness arose in data on postprandial BA flow within the small intestine, measured in seven ^26^, three ^27^, or a single individual ^25^. Ethical constraints prevented organ sampling from healthy subjects, requiring data from patients undergoing unrelated surgeries. While this approach provided the only means to inform BA concentration outside systemic blood circulation, questions lingered about its accuracy.

Moreover, only two studies involving multiple meals were identified, and these could not be harmonized with numerous single meal studies. Dirksen et al.’s study ^29^ reported postprandial responses for three meals, with the first meal showing no notable increase in blood BA levels, possibly attributed to a systematic measurement error. The second and third responses exhibited more reasonable dynamics, albeit with notably high reported levels. Conversely, the five meal study by Al-Khaifi et al. ^28^ reported consistent but low BA levels.

This discrepancy was in part attributed to the fact that BAs exhibit notable endogenous variability among individuals ^32-34^. This variation is evident in both fasting and postprandial states, with the latter showcasing pronounced variability. To address this complexity, a meta-analysis of various studies was conducted to estimate variations between studies and within the population. Additionally, a detailed exploration of BA metabolism variability was achieved through population simulation, a robust method for understanding complex biological and physiological systems. This simulation facilitated close alignment of the models with distribution patterns in reported data, particularly in the context of postprandial responses in the bloodstream. These outcomes collectively provide compelling evidence of the models’ quality, instilling confidence in its suitability for subsequent analyses and predictions.

Due to its mechanistic framework, the model has potential for extrapolation to diverse conditions, including diseases.

Unlike previous studies {de Bruijn, 2022 #62;De Bruijn, 2024 #63;Guiastrennec, 2018 #48;Sips, 2018 #46;Voronova, 2020 #47;Woodhead, 2014 #61}, the previously published models were not constructed upon existing knowledge; consequently, physiological parameters had to be established anew. In this study, all model parameters are linked to specific physiological functions, avoiding a mere empirical description of processes. This characteristic allows for extrapolation to novel scenarios. As a result, the models introduced in this study have the capacity to elucidate bile acid levels throughout the entire body, making them particularly adept for predicting outcomes in clinically relevant contexts.

In subsequent analyses, refinements may be applied to enhance the structural aspects of the physiologically-based model, addressing inherent limitations. The current version of the computational model focuses solely on total levels of the predominant BA species, omitting conjugation status and less common secondary BAs. Consequently, the models fall short of capturing the complete complexities of the intricate BA pool, potentially introducing a systemic bias in predictions due to variations in kinetics among different BA species ^42^.

The primary limitation of this work arises from the limited available data, especially concerning BA levels and microbial density in the gut. Therefore, any predictions related to intestinal BAs should be approached with utmost caution. Despite these constraints, the models effectively reproduced BA composition and levels throughout the body, precisely capturing inter-individual variability in postprandial responses. We hence expect to support alterations in bile acid levels in health and disease in the future.-

## Acknowledgments

Funding was provided from the German Research Foundation (DFG) – Project-ID 403224013 – “SFB 1382”.

## Author Contributions

Bastian Kister: Methodology, Software, Validation, Formal analysis, Writing--original draft, Writing--review & editing, Visualization

Lars M. Blank: Writing--review & editing

Maike Pollmanns: Writing--review & editing

Theresa H. Wirtz: Conceptualization, Resources, Writing--original draft, Writing--review & editing, Supervision

Lars Kuepfer: Conceptualization, Resources, Writing--original draft, Writing--review & editing, Supervision

## Declaration of interest

The authors have no conflicts of interest to declare

## Methods

### Materials Availability

This study did not generate new unique reagents.

### Data and code availability

The computational model file is available on http://www.ukaachen.de/kuepfer.

Any additional information required to reanalyze the data reported in this paper is available from the lead contact upon request.

### Computational methods and PBPK modelling

Physiologically-based pharmacokinetic (PBPK) models describe the physiology of an organism at a large level of detail. Organs are explicitly represented in PBPK models and they are linked through systemic vascular circulation. Tissue concentrations can be simulated in PBPK models, even if they are experimentally inaccessible. Parameters in PBPK models explicitly represent specific physiological functions and they are taken from previously curated collections of parameters including organ volumes, surface areas, tissue composition and blood perfusion rates, respectively. For that reason, identification of PBPK models is limited to very few parameters, usually related to active processes underlying compound distribution of as well as elimination. PBPK models are hence based on a large amount of prior knowledge including detailed description of physiological processes such as enterohepatic circulation or absorption in the intestine

#### Kinetic rate laws

Describing BA synthesis, a constant flux within the intracellular space of the liver was assumed. The following rate law was applied :

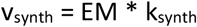

where EM represents the amount of catalyzing enzyme and k_synth_ the absolute synthesis rate.

For the remaining enzymatic reactions and transport processes, Michaelis-Menten kinetics were applied, following in mice:

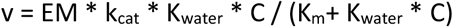

with K_water_ describing the partition coefficient of the BA between water and the source compartment of the BA, K_m_ is the Michaelis-Menten constant and C the BA concentration in the source compartment. k_cat_ represents the number of substrate BA each enzyme site converts to product per unit time, and in which the enzyme is working at maximum efficiency and is calculated as

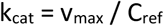

with v_max_ being the maximum rate of reaction and C_ref_ the enzyme reference concentration of 1 µM. For renal excretion, tubular excretion with Michaelis-Menten kinetic was selected within PKSim:

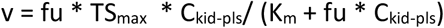

with TSmax describes the intrinsic maximum rate for tubular secretion, fu the fraction unbound BA in blood plasma and C_kid-pls_ the BA concentration within the plasma sub-compartment of the kidney.

### Calculations

Parameter fitting was performed with the Monte Carlo algorithm implemented in the Open Systems Pharmacology suite. Residual calculation was set to linear and weights were derived from measured SD.

Sensitivity analysis was performed using the default settings within MoBi®. For sensitivity coefficients a threshold of 1 and -1 was applied.

Most reactions were defined as simple Michaelis-Menten kinetics, for BA synthesis a constant flux was assumed.

### Quantification and statistical analysis

The physiologically-based model of bile acid metabolism was established in PK-Sim® and further reactions and adjustments were done in MoBi® (Open Systems Pharmacology suite Version 11.150). Model simulations were performed using the ospsuite-R package in R (version 11.0.123).

Plotting and statistical testing was done with custom Python scripts.

Based on the literature dataset and subsequent meta-analysis, it is possible to provide a more detailed characterization of the ‘healthy’ reference state for BA metabolism:

- Systemic circulation The baseline BA levels in systemic circulation were found to be approximately 2.83 µM, with CDCA being the most abundant BA species. Conjugated BAs accounted for about 61% of the total, with glycine-conjugates making up approximately 87% of this fraction. Following food intake, single-meal studies reported an increase of up to 4.3-fold, while multiple-meal studies showed increases of up to 6.3-fold (Supplementary Table}).
- Liver and portal blood Concentrations in portal blood were reported for CA, CDCA, and DCA, with primary BAs being nearly twice as abundant as secondary BAs. CA and CDCA together comprised about 42% of the total BA pool, approximately 61 µM. Unfortunately, information on the conjugation status of BAs in these organs could not be obtained (Supplementary Tables and}). In all three organs (systemic circulation, portal blood, and liver), DCA was the most abundant secondary BA, while LCA and UDCA were equally abundant.
- Biliary composition Biliary composition of BAs exhibited high variability, but CAs, CDCAs, and DCAs each comprised approximately 30% of the BA pool in bile. Glycine-conjugated BAs were more abundant than T-BAs, and uBAs, as well as LCAs and UDCAs, together accounted for less than 3% (Supplementary Table).
- {Intestinal BA flow Basal intestinal BA flow through the SI was highest in the duodenum, at 7.44 µmol/min, and decreased along the gut axis (jejunum: 5.29 µmol/min; ileum: 2.66 µmol/min). In contrast, the most significant postprandial increase was observed in the ileum and diminished towards the duodenum (ileum: 40-fold; jejunum: 22-fold; duodenum: 9-fold). See also supplementary tables

Schaap, F. G., Trauner, M. & Jansen, P. L. Bile acid receptors as targets for drug development. *Nat Rev Gastroenterol Hepatol* **11**, 55-67 (2014). https://doi.org/10.1038/nrgastro.2013.151

## Notes

### Competing Interest Statement

The authors have declared no competing interest.

