## Supplementary Information for "A physiologically-based model of bile acid metabolism in humans"

### Supporting information

Table 1: Data of bile acid concentration in venous blood plasma derived from literature used for model development.

| Bile acid | Value | SEM | Unit | Number of studies | Source |
|---|---|---|---|---|---|
| tBA | 2.83 | 1.1 | μM | 18 | [326; 327; 328; 330; 331; 354; 356; 357; 359; 360; 361; 362; 363; 364; 365; 366; 367; 368] |
| Conj-BA | 1.74 | 0.72 | μM | 4 | [327; 356; 362; 367] |
| Glyco-BA | 1.52 | 0.66 | μM | 3 | [356; 362; 367] |
| tCA | 0.50 | 0.25 | μM | 5 | [327; 354; 361; 362; 367] |
| G-CA | 0.18 | 0.14 | μM | 2 | [362; 367] |
| T-CA | 0.06 | 0.05 | μM | 2 | [362; 367] |
| CA | 0.23 | 0.15 | μM | 2 | [362; 367] |
| tCDCA | 1.21 | 0.53 | μM | 5 | [327; 354; 361; 362; 367] |
| G-CDCA | 0.76 | 0.43 | μM | 2 | [362; 367] |
| T-CDCA | 0.13 | 0.29 | μM | 2 | [362; 367] |
| CDCA | 0.30 | 0.20 | μM | 2 | [362; 367] |
| tDCA | 0.82 | 0.39 | μM | 5 | [327; 354; 361; 362; 367] |
| G-DCA | 0.34 | 0.22 | μM | 2 | [362; 367] |
| T-DCA | 0.07 | 0.06 | μM | 2 | [362; 367] |
| DCA | 0.44 | 0.28 | μM | 2 | [362; 367] |
| tLCA | 0.26 | 0.18 | μM | 3 | [361; 362; 367] |
| G-LCA | 0.07 | 0.06 | μM | 2 | [362; 367] |
| T-LCA | 0.04 | 0.04 | μM | 2 | [362; 367] |
| LCA | 0.02 | 0.01 | μM | 1 | [367] |
| tUDCA | 0.20 | 0.11 | μM | 4 | [327; 359; 362; 367] |
| G-UDCA | 0.15 | 0.12 | μM | 2 | [362; 367] |
| T-UDCA | 0.01 | 2e-3 | μM | 1 | [367] |
| UDCA | 0.09 | 0.08 | μM | 2 | [362; 367] |

Table 2: Data of bile acid concentration in portal blood plasma derived from literature used for model development.

| Bile acid | Value | SEM | Unit | Number of studies | Source |
|---|---|---|---|---|---|
| tCA | 5.16 | 0.71 | μM | 4 | [328; 329; 330; 331] |
| tCDCA | 5.32 | 1.48 | μM | 4 | [328; 329; 330; 331] |
| tDCA | 3.62 | 1.04 | μM | 4 | [328; 329; 330; 331] |

Table 3: Data of bile acid pool size in liver derived from literature used for model development.

| Bile acid | Value | SEM | Unit | Number of studies | Source |
|---|---|---|---|---|---|
| tCA | 60.99 | 4.18 | μmol | 4 | [9; 332; 333; 334] |
| tCDCA | 62.48 | 3.99 | μmol | 4 | [9; 332; 333; 334] |
| tDCA | 14.29 | 8.77 | μmol | 4 | [9; 332; 333; 334] |
| tLCA | 2.16 | 1.59 | μmol | 4 | [9; 332; 333; 334] |
| tUDCA | 5.01 | 1.24 | μmol | 4 | [9; 332; 333; 334] |

Table 4: Data of synthesis and excretion rates derived from literature used for model development.

| Rates | Value | SEM | Unit | Number of studies | Source |
|---|---|---|---|---|---|
| CA synthesis | 0.62 | 0.20 | mmol/day | 5 | [325; 369; 370; 371; 372] |
| CDCA synthesis | 0.37 | 0.10 | mmol/day | 4 | [369; 370; 371; 372] |
| Renal excretion | 4.29 | 3.1 | μmol/day | 4 | [363; 373; 374; 375] |

Table 5: Data of bile acid composition in feces derived from literature used for model development.

| Bile acid | Value | SEM | Unit | Number of studies | Source |
|---|---|---|---|---|---|
| tCA | 3.5 | 2.8 | % | 2 | [362; 376] |
| tCDCA | 3.4 | 2.7 | % | 2 | [362; 376] |
| tDCA | 58.2 | 16.1 | % | 2 | [362; 376] |
| tLCA | 35.0 | 15.2 | % | 2 | [362; 376] |
| tUDCA | 1.4 | 1.2 | % | 2 | [362; 376] |

Table 6: Data of bile acid composition in bile derived from literature used for model development.

| Bile acid | Value | SEM | Unit | Number of studies | Source |
|---|---|---|---|---|---|
| CA | 0.09 | 0.07 | % | 1 | [377] |
| G-CA | 29.77 | 58.26 | % | 1 | [377] |
| T-CA | 2.45 | 3.49 | % | 1 | [377] |
| G-CDCA | 20.51 | 43.56 | % | 1 | [377] |
| T-CDCA | 8.20 | 14.61 | % | 1 | [377] |
| G-DCA | 27.61 | 64.83 | % | 1 | [377] |
| T-DCA | 8.56 | 16.45 | % | 1 | [377] |
| G-LCA | 0.55 | 1.30 | % | 1 | [377] |
| T-LCA | 0.22 | 0.57 | % | 1 | [377] |
| UDCA | 0.00 | 0.00 | % | 1 | [377] |
| G-UDCA | 1.97 | 2.65 | % | 1 | [377] |
| T-UDCA | 0.08 | 0.25 | % | 1 | [377] |

Table 7. Data of intestinal bile flows derived from literature used for model development.

| Flows | Value | SEM | Unit | Number of studies | Source |
|---|---|---|---|---|---|
| Duodenal bile flow (basal) | 7.44 | 3.5 | µmol/min | 1 | [323] |
| Jejunal bile flow (basal) | 5.29 | 2.16 | µmol/min | 1 | [323] |
| Ileal bile flow (basal) | 2.66 | 1.65 | µmol/min | 1 | [323] |
| Per day | 30.0 | 3.40 | mmol/day | 1 | [324] |
| First h postprandial | 2.56 | 0.37 | mmol/h | 1 | [324] |
| Overnight (12h) | 0.90 | 0.11 | µmol/min | 1 | [324] |

Table 8. Literature data of postprandial responses in small intestine used for model development. Error was calculated using all measurements within one segment as individual measurements as no measurement error was determined within the study.

| Segment | Time [min] | Flux [µmol/min] | Error (calculated) | Source |
|---|---|---|---|---|
| Duodenum | 30 | 59.84 | 17.00 | [323] |
| Duodenum | 60 | 64.88 | 17.00 | [323] |
| Duodenum | 90 | 63.54 | 17.00 | [323] |
| Duodenum | 120 | 48.37 | 17.00 | [323] |
| Duodenum | 150 | 41.87 | 17.00 | [323] |
| Duodenum | 180 | 43.56 | 17.00 | [323] |
| Duodenum | 210 | 26.75 | 17.00 | [323] |
| Duodenum | 240 | 18.29 | 17.00 | [323] |
| Jejunum | 30 | 33.40 | 32.96 | [323] |
| Jejunum | 60 | 117.52 | 32.96 | [323] |
| Jejunum | 90 | 54.78 | 32.96 | [323] |
| Jejunum | 120 | 26.35 | 32.96 | [323] |
| Jejunum | 150 | 31.97 | 32.96 | [323] |
| Jejunum | 180 | 26.05 | 32.96 | [323] |
| Jejunum | 210 | 22.33 | 32.96 | [323] |
| Jejunum | 240 | 15.84 | 32.96 | [323] |
| Ileum | 30 | 16.04 | 39.59 | [323] |
| Ileum | 60 | 105.49 | 39.59 | [323] |
| Ileum | 120 | 82.42 | 39.59 | [323] |
| Ileum | 180 | 31.86 | 39.59 | [323] |
| Ileum | 210 | 17.91 | 39.59 | [323] |
| Ileum | 240 | 12.81 | 39.59 | [323] |

Table 9: Fitted parameter values of enzymatic reactions and postprandial responses in the BA model.

| Parameter | Value | Unit |
|---|---|---|
| Parameter | Value | Unit |
| Synthesis rate (tCA) | 9.5E-01 | µmol/min |
| Synthesis rate (tCDCA) | 4.4E-01 | µmol/min |
| Renal Clearance (Km) | 9.5E+02 | µmol/l |
| Renal Clearance (TSmax) | 6.0E-03 | µmol/l/min |
| tCA →tDCA (Km) | 2.4E+01 | µmol/l |
| tCA →tDCA (vmax) | 2.0E+02 | µmol/l/min |
| tCDCA →tLCA (Km) | 1.1E+02 | µmol/l |
| tCDCA →tLCA (vmax) | 9.5E+02 | µmol/l/min |
| tCDCA →tUDCA (Km) | 1.0E+03 | µmol/l |
| tCDCA →tUDCA (vmax) | 4.3E+03 | µmol/l/min |
| tUDCA →tLCA (Km) | 9.5E+02 | µmol/l |
| tUDCA →tLCA (vmax) | 1.8E+01 | µmol/l/min |
| Gallbladder emptying lag time | 9.00 | min |
| Gallbladder ejection half-time | 59.73 | min |
| EHC continuous fraction | 0.50 | |
| Gallbladder ejection fraction | 0.40 | |

Table 10: Fitted parameter values of hepatic transport reactions in the BA model.

| Parameter | Value | Unit |
|---|---|---|
| BSEP (tCA; Km) | 9.5E+02 | µmol/l |
| BSEP (tCDCA; Km) | 9.6E+02 | µmol/l |
| BSEP (tDCA; Km) | 2.9E+02 | µmol/l |
| BSEP (tLCA; Km) | 9.1E+02 | µmol/l |
| BSEP (tUDCA; Km) | 9.9E+02 | µmol/l |
| BSEP (tCA; vmax) | 9.2E+00 | µmol/l/min |
| BSEP (tCDCA; vmax) | 6.5E+00 | µmol/l/min |
| BSEP (tDCA; vmax) | 1.4E+01 | µmol/l/min |
| BSEP (tLCA; vmax) | 4.4E+01 | µmol/l/min |
| BSEP (tUDCA; vmax) | 5.3E+00 | µmol/l/min |
| NTCP (tCA; Km) | 7.6E+02 | µmol/l |
| NTCP (tCDCA; Km) | 9.2E+02 | µmol/l |
| NTCP (tDCA; Km) | 4.5E+01 | µmol/l |
| NTCP (tLCA; Km) | 8.0E+02 | µmol/l |
| NTCP (tUDCA; Km) | 6.3E+02 | µmol/l |
| NTCP (tCA; vmax) | 2.4E+03 | µmol/l/min |
| NTCP (tCDCA; vmax) | 4.8E+02 | µmol/l/min |
| NTCP (tDCA; vmax) | 4.9E+03 | µmol/l/min |
| NTCP (tLCA; vmax) | 2.6E+03 | µmol/l/min |
| NTCP (tUDCA; vmax) | 1.1E+03 | µmol/l/min |

Table 11: Fitted parameter values of intestinal transport reactions in the BA model

| Parameter | Value | Unit |
|---|---|---|
| ASBT (tCA; Km) | 1.0E+03 | $\mu$mol/l |
| ASBT (tCDCA; Km) | 1.6E+02 | $\mu$mol/l |
| ASBT (tDCA; Km) | 1.3E+02 | $\mu$mol/l |
| ASBT (tLCA; Km) | 2.4E+01 | $\mu$mol/l |
| ASBT (tUDCA; Km) | 4.4E+01 | $\mu$mol/l |
| ASBT (tCA; vmax) | 4.1E+02 | $\mu$mol/l/min |
| ASBT (tCDCA; vmax) | 1.3E+01 | $\mu$mol/l/min |
| ASBT (tDCA; vmax) | 5.0E+03 | $\mu$mol/l/min |
| ASBT (tLCA; vmax) | 2.1E+03 | $\mu$mol/l/min |
| ASBT (tUDCA; vmax) | 3.8E+03 | $\mu$mol/l/min |
| OST$\alpha/\beta$ (tCA; Km) | 9.2E-01 | $\mu$mol/l |
| OST$\alpha/\beta$ (tCDCA; Km) | 1.0E+03 | $\mu$mol/l |
| OST$\alpha/\beta$ (tDCA; Km) | 6.7E+01 | $\mu$mol/l |
| OST$\alpha/\beta$ (tLCA; Km) | 9.6E+02 | $\mu$mol/l |
| OST$\alpha/\beta$ (tUDCA; Km) | 2.2E+01 | $\mu$mol/l |
| OST$\alpha/\beta$ (tCA; vmax) | 5.0E+03 | $\mu$mol/l/min |
| OST$\alpha/\beta$ (tCDCA; vmax) | 1.3E+00 | $\mu$mol/l/min |
| OST$\alpha/\beta$ (tDCA; vmax) | 5.0E+03 | $\mu$mol/l/min |
| OST$\alpha/\beta$ (tLCA; vmax) | 2.5E+00 | $\mu$mol/l/min |
| OST$\alpha/\beta$ (tUDCA; vmax) | 5.0E+03 | $\mu$mol/l/min |

Table 12: Fitted parameter values of enzymatic reactions and postprandial responses in the BA model.

| Liver conc. [$\mu$M] | Parameter PK | Model | Sensitivity coefficient |
|---|---|---|---|
| tDCA | C_tEnd | pH (intracellular) | 31.9944845 |
| tCA | C_tEnd | pH (intracellular) | 17.11378011 |
| tUDCA | C_tEnd | pH (intracellular) | 13.97256935 |
| tCA | C_max | pH (intracellular) | 12.07650294 |
| tCDCA | C_tEnd | pH (intracellular) | 5.85872439 |
| tCDCA | C_max | pH (intracellular) | 4.991866588 |
| tUDCA | C_max | LU-ColonTransversum-pH | 4.505388517 |
| tUDCA | C_tEnd | LU-ColonTransversum-pH | 4.240910746 |
| tLCA | C_tEnd | pH (intracellular) | 4.073137463 |
| tLCA | C_max | pH (intracellular) | 4.019450905 |
| tUDCA | C_tEnd | LU-ColonAscendens-pH | 3.882990266 |

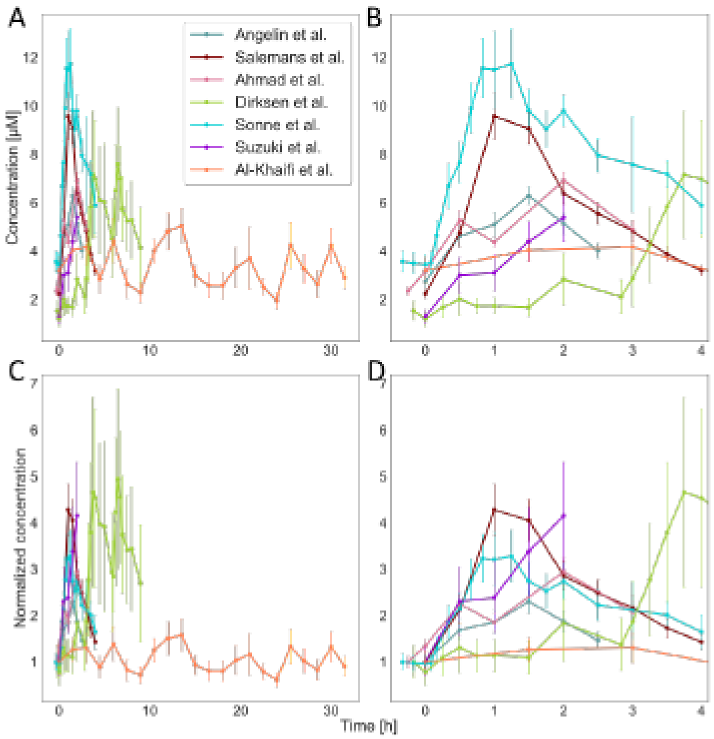


Figure 1: Data of postprandial responses in venous blood. Postprandial responses in venous blood plasma used for model development. Displayed are the profiles as taken from literature (A), only for the first meal response (B) and corresponding profiles normalized to the initial time point (C and D). Errors bars represent the SD. Data was taken from [326; 327; 354; 355; 356; 357; 358]
